# Transcriptomic analysis of naïve human embryonic stem cells cultured in three-dimensional PEG scaffolds

**DOI:** 10.1101/2020.09.16.299909

**Authors:** Christina McKee, Christina Brown, Shreeya Bakshi, Keegan Walker, Chhabi K. Govind, G. Rasul Chaudhry

## Abstract

Derivation of primed and naïve human embryonic stem cells (ESCs) have prompted an increased interest in devising culture conditions for maintaining their pluripotency and differential potential. Naïve ESCs are characterized by improved viability, proliferation, and differentiation capacity in comparison to primed ESCs. However, traditional two-dimensional (2-D) cell culture techniques fail to mimic the three-dimensional (3-D) *in vivo* microenvironment, which results in altered morphological and molecular characteristics of ESCs. Here, we describe the use of 3-D self-assembling scaffolds that support growth and maintenance of the naïve state characteristics of human ESC line, Elf1. Scaffolds were formed via a Michael addition reaction upon combination of two 8-arm polyethylene glycol (PEG) polymers functionalized with thiol (PEG-8-SH) and acrylate (PEG-8-Acr) end groups. 3-D scaffolds not only maintained the naïve state, but also supported long-term growth for up to 3 weeks without requiring routine passaging and manipulation. 3-D grown cells exhibited upregulation of core (*OCT4*, *NANOG*, and *SOX2*) and naïve (*KLF17*, *KLF4*, *TFCP2L1*, *DPPA3*, and *DNMT3L*) genes. These genes returned to normal levels when 3-D grown cells were propagated under 2-D culture conditions. Examination of RNA-sequencing demonstrated significant changes in gene expression profiles between 2-D and 3-D grown Elf1 cells. Gene Ontology analysis revealed upregulation of biological processes involved in the regulation of transcription and translation, as well as β-catenin-TCF complex assembly, extracellular matrix organization, and chromatin remodeling in 3-D grown Elf1 cells. 3-D culture conditions also induced upregulation of genes associated with several signaling pathways including Wnt signaling and focal adhesion. However, p53 signaling pathway associated genes were downregulated under these culture conditions. Our findings provide insight into the possible mechanisms of prolonged self-renewal as well as upregulation of pluripotent genes stimulated by the transduction of mechanical signals from the 3-D microenvironment.

## 1. Introduction

Embryonic stem cells (ESCs) are pluripotent cells capable of unlimited self-renewal and differentiation into all three germ layers [1, 2], making them an attractive source for the development of *in vitro* disease models and drug discovery as well as tissue engineering and cellular therapies [3–5]. These applications require stringent conditions for characterization and expansion as well as differentiation of cells into homogenous populations of progenitors or derivatives. However, these processes are hindered both by limited understanding of pluripotent cell biology as well as current culture techniques [6, 7].

Traditionally, human ESCs are derived from the inner cell mass of a preimplantation blastocyst, exhibiting growth characteristics similar to mouse epiblast stem cells (EpiSCs) *in vitro* [8]. In contrast, mouse ESCs, isolated at an earlier developmental stage, show improved clonal growth and viability following single cell dissociation and are capable of contributing to blastocyst chimeras [9]. Based on their divergent characteristics *in vitro*, two distinct stages of pluripotency were proposed, an early naïve state (mouse ESCs) and a primed state (human ESCs and mouse EpiSCs) [10].

Naïve pluripotency affords certain advantages over traditionally derived primed ESCs, including improved proliferation and developmental potential as well as single-cell cloning efficiency and genome editing [11, 12]. Consequently, a great deal of effort has been focused on both the reprogramming and establishment of naïve human ESC lines [9, 13, 14].

Several reports described successful culture of human ESCs in a naïve pluripotent state using ectopic expression of select genes and inhibitor cocktails [11, 15–19]. Recently, improved culture conditions led to derivation of naïve human ESC lines directly from six to eight-cell embryos [20–22]. This has led to an increased understanding of the importance of defined culture media conditions for the modulation of cell signaling and maintenance of pluripotency *in vitro*.

However, ESC culture practices are still antiquated, relying on adherence to two-dimensional (2-D) tissue-culture polystyrene plates often coated with natural or synthetic substrates mimicking extracellular matrix (ECM) components for the propagation of cells [6]. Generally, 2-D culture conditions fail to mimic the native three-dimensional (3-D) microenvironments that are responsible for the regulation of cell fate *in vivo*, allowing for dynamic spatial interactions between cells, ECM components, and gradients of soluble factors via biochemical, mechanical, and structural stimuli [23–25]. 2-D culture fails to accurately reproduce the physiology of the embryo [26] resulting in the spontaneous differentiation of ESC colonies *in vitro* [27, 28].

Therefore, development of 3-D culture techniques via the incorporation of bioinductive materials has gained traction in recent years [6]. Natural biomaterials such as collagen type 1, gelatin, alginate, and hyaluronic acid have been studied for the 3-D propagation of human ESCs; [29–33]. However, these materials have been shown to induce biological signaling and variability in culture [34]. In contrast, inert synthetic biomaterials can be more easily modified, allowing for tunable mechanical and biophysical properties as well as minimizing batch-to-batch variation [35]. Therefore, synthetic biomaterials may prove to be ideal for the development of the defined 3-D culture conditions needed for the propagation of human ESCs. We have previously demonstrated 3-D culture of naïve mouse ESCs [36] and primed human ESCs [37] using hydrogel scaffolds that resulted in modulation of gene expression, and maintenance of pluripotent cells. In this study, we investigated 3-D culture of naïve human ESCs (Elf1 cells) using self-assembling scaffolds comprised of two synthetic 8-arm polyethylene glycol (PEG) polymers functionalized with thiol (PEG-8-SH) and acrylate end groups (PEG-8-Acr). 3-D culture in the PEG-8-SH/PEG-8-Acr scaffolds supported ESC self-renewal and pluripotency. To determine the effect of culture in the PEG-8-SH/PEG-8-Acr scaffolds on the maintenance of pluripotency, we compared changes in the transcriptomic profile of naïve ESCs grown in 2-D and 3-D culture conditions. The improved understanding of the molecular functions and pathways associated with continued self-renewal of naïve ESCs enriched in 3-D culture may lead to improvements in *in vitro* culture techniques, thus promoting their use in basic and applied applications.

## 2. Materials and Methods

### 2.1 Maintenance of naïve ESCs in 2-D culture

Elf1 cells, a naïve human ESC line isolated from an 8-cell human embryo, were procured from Wicell (Madison, WI), and maintained according to a previously published protocol [22]. Briefly, Elf1 cells were grown adherent to mouse embryonic feeder (MEF) coated 6-well plates, and incubated in medium containing KnockOut/F12 DMEM with GlutaMax (Life Technologies, Carlsbad, CA) with 20% KnockOut serum replacement (Life Technologies), 1mM sodium pyruvate (Life Technologies), 0.1mM 2-Mercaptoehtanol (Life Technologies), 1% non-essential amino acids (Life Technologies), and 0.2% penicillin-streptomycin solution (Life Technologies), and supplemented with 12ng/ml FGF2 (Prospec, Ness Ziona, Israel), 1.5μM CHIR99021 (Cayman Chemical, Ann Arbor, MI), 0.4μM PD03296501 (Caymen Chemical) and 0.01μg/ml human LIF (Prospec). Prior to encapsulation in 3-D scaffolds, cells were transitioned to feeder-free culture in Matrigel-coated 6-well plates (Fisher Scientific, Pittsburgh, PA).

### 2.2 Self-assembly of 3-D scaffolds via thiol-Michael addition reaction for encapsulation of cells

8-arm PEG polymers functionalized with thiol and acrylate end groups were purchased from JenKem Technology USA (Plano, TX), and stored in the dark at −20°C. PEG-8-SH and PEG-8-Acr polymers were combined to facilitate self-assembly, as previously described [37]. Briefly, PEG-8-SH and PEG-8-Acr were separately dissolved in culture media at 2.5 w/v% polymer concentration and mixed thoroughly at a 1:1 molar ratio in the presence of oxygen for polymerization via a thiol-Michael addition reaction. The schematic of scaffold formation is depicted in Figure 1. For 3-D culture, ESCs were dissociated into single cells using Accutase (Life Technologies), pelleted at 5×10^6^ cells/ml, resuspended in the PEG-8-SH/PEG-8-Acr polymer mixture, and transferred to syringe molds for polymerization, forming 100μl scaffolds. Following scaffold formation, scaffolds were placed in 24-well culture plates (Fisher Scientific), supplemented with culture medium, and incubated at 37°C and 5% CO_2_. The medium was changed daily or as needed.

**Figure 1.**
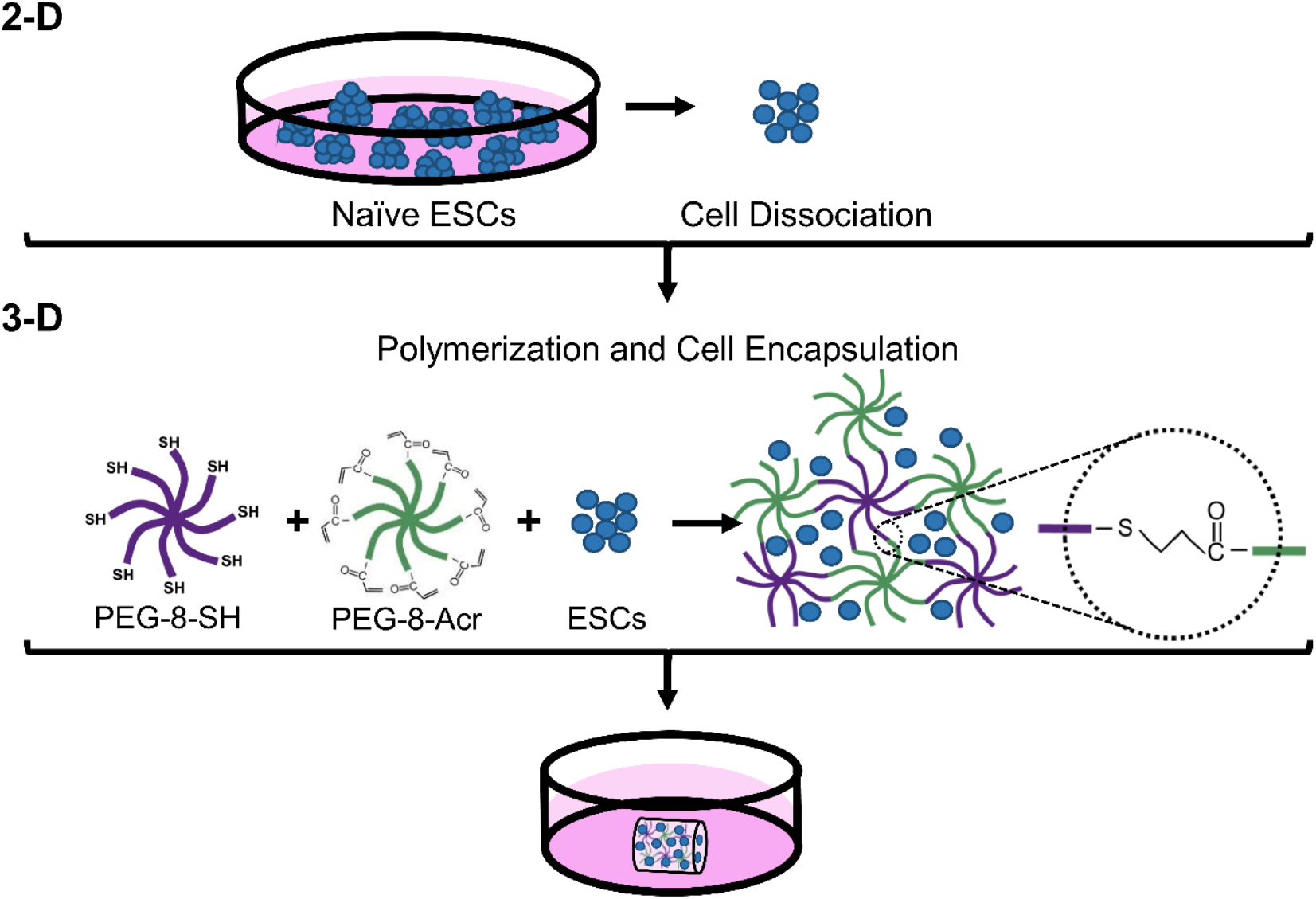
Encapsulation and 3-D culture of naïve ESCs in self-assembling scaffolds. Elf1 cells grown in 2-D culture conditions were dissociated into single cells, and mixed with self-assembling end-functionalized polymers, PEG-8-SH and PEG-8-Acr. The polymer/cell mixture was then transferred to a syringe mold and allowed to form 3-D scaffolds, encapsulating the cells. The scaffolds were then transferred to 24-well plates containing culture medium.

### 2.3 Cell proliferation of 3-D grown ESCs

Proliferation of cells grown under 3-D culture conditions was monitored by phase-contrast microscopy and quantified at various time points using an MTT proliferation assay. Samples were prepared in triplicate experiments and treated with 5mg/ml MTT reagent (Sigma, St. Louis, MO, USA), incubated at 37°C for 4 hrs, and subsequently treated with isopropanol/HCL (15:1) to solubilize the formazan product produced by viable cells [38]. The absorbance of the solubilized formazan was then measured at 570nm using an Epoch Microplate Spectrophotometer (BioTek, Winooski, VT), and the background absorbance was subtracted from all experimental values.

### 2.4 Teratoma formation assay

Pluripotency of 3-D grown cells was assessed by teratoma formation assays performed in triple experiments. ESCs (1×10^6^) were dissociated into single cells following Accutase treatment, resuspended in PBS, mixed with an equal volume of Matrigel (BD Biosciences, San Jose, CA), and injected subcutaneously into the flanks of 4-week-old immunocompromised Fox Chase Severe Combined Immunodeficiency (SCID) Beige mice (Charles River, Wilmington, MA) using a Hamilton syringe. Animals were monitored daily and humanely euthanized by CO_2_ overdose following teratoma formation at 10-12 weeks post-injection. Explanted teratomas were either fixed for histological analysis or flash frozen for RNA isolation and assessed by immunohistochemistry and quantitative real time-polymerase chain reaction (qRT-PCR) for germ layer marker expression, respectively. All procedures involving animals were approved by the Institutional Animal Care and Use Committee of Oakland University (IACUC protocol number: 17031).

### 2.5 Gene expression analysis using qRT-PCR

Total cellular mRNA was isolated from cells grown in 2-D and 3-D culture conditions using the GeneJET RNA purification kit (Thermo Fisher Scientific). For analysis of germ layer markers, total RNA was extracted from teratoma tissue (100-250mg) using RNeasy Midi kit (Qiagen, Germantown, MD) per the manufacturer’s instructions. cDNA was synthesized using the iScript kit (BioRad, Hercules, CA). qRT-PCR was performed using SsoAdvanced SYBR Green Supermix (Bio-Rad) and the CFX96 Real-Time PCR system. Triplicate reactions were normalized to reference genes, *HMBS* and *GAPDH*. Primers (IDT Technologies, Coralville, IA) used in this study are presented in Table 1.

**Table 1.**
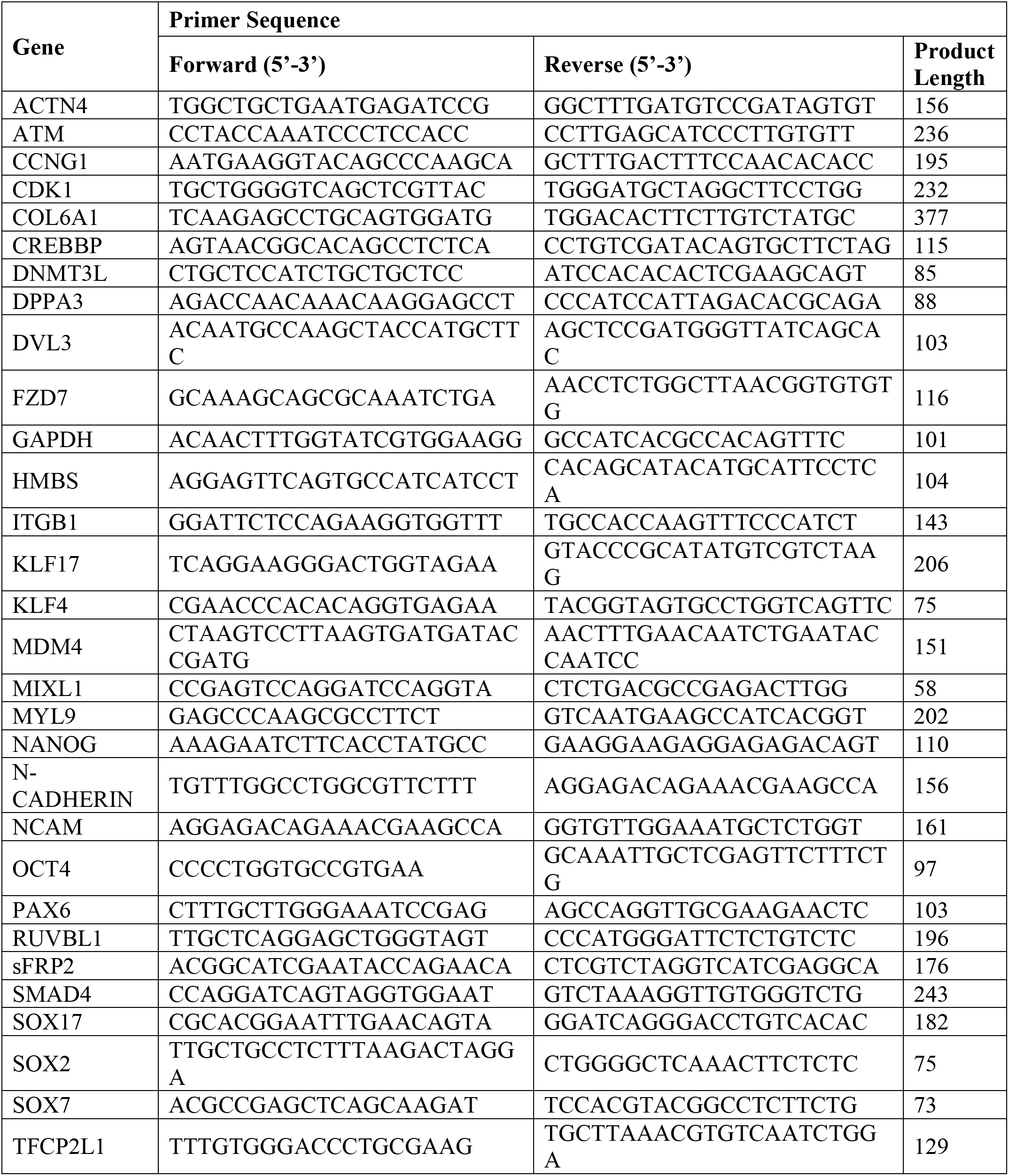
List of human primer sequences used in qRT-PCR.

### 2.6 RNA-sequencing (RNA-seq)

Total RNA from 2-D and 3-D grown cells was isolated as described. RNA was quantified and qualified using an Agilent2100 Bioanalyzer (Agilent Technologies, Palo Alto, CA) and Qubit Assay (Life Technologies). RNA with an RNA integrity number (RIN) of 10 was used as input material for library preparation. cDNA libraries were prepared using the KAPA RNA HyperPrep Kit with RiboErase (HMR) (Kapa Biosystems, Wilmington, MA) according to the manufacturer’s protocol [39], and sent for RNA-seq transcriptome analysis.

GENEWIZ (South Plainfield, NJ) performed 2 × 150bp paired end read sequencing on the Illumina NovaSeq/HiSeq. An average of 47 million reads were obtained for each sample. Fragments were mapped to reference human genome assembly hg38 (with an average mapping efficiency of 94%) and differential gene expression analysis was performed using the galaxy platform (https://usegalaxy.org/) [40] as detailed below. RNA-Seq analyses were performed on two independent biological replicates.

### 2.7 Bioinformatic analysis

The quality of the raw reads were assessed using FastQC toolkit v0.72 [41]. Low quality reads and adapters were trimmed using Trim Galore! v0.6.3 [42]. HiSat2 v2.1.0 [43] was then used for mapping the raw reads to the reference genome human genome assembly hg38. The expression for each gene were evaluated using featureCounts v1.6.4 [44] based on annotation from GENCODE release 30 [45]. The raw counts were then normalized and differential gene expression was determined using DESeq2 v2.11.40.6 [46]. Genes with an adjusted p-value or Benjamini-Hochberg false discovery rate (FDR) adjustment < 0.05 were considered significant unless otherwise specified.

Biological significance and functional assessment of genes differentially expressed between 2-D and 3-D culture conditions were analyzed by Gene Ontology (GO) assessment. Protein ANalysis THrough Evolutionary Relationships (PANTHER) [47] and the Kyoto Encyclopedia of Genes and Genomes (KEGG) pathway [48] databases were used to analyze biological pathways associated with differentially expressed genes (DEGs). In addition, enrichment of DEGs associated with cellular compartment, molecular function and biological processes was also performed using Enrichr [49, 50]. Significance was defined by the recommended p-value <0.05. Heatmaps were generated using Heatmapper (http://www.heatmapper.ca/) [51].

### 2.8 Immunohistochemical analysis

Protein expression of germ layer markers was investigated by fluorescent immunohistochemical staining. Teratomas were fixed overnight in 4% paraformaldehyde and embedded in O.C.T solution for cryosectioning into 10*μ*m sections. For immunohistochemical analysis, cryosections were permeabilized with 0.5% Triton X-100 (Sigma) for 10 mins and blocked with 2% BSA (Sigma) for 1hr at room temperature. Samples were incubated overnight at 4°C with primary antibodies (1:100 dilution) against GATA4 (sc-25310, Santa Cruz Biotechnology, Santa Cruz, CA), BRACHYURY (sc-20109, Santa Cruz), and TUJ1 (sc-58888, Santa Cruz) representing endoderm, mesoderm, and ectoderm germ layers, respectively. Primary antibody treated samples were then washed with PBS, stained with anti-rabbit Alexa Fluor 488 (A32731, Thermo Fisher Scientific) or anti-mouse Cy3-labeled IgG (072-01-18-06, KPL, Gaithersburg, MD, United States) secondary antibodies at 1:200 dilution for 2 hrs at room temperature, and counterstained with DAPI (1mg/mL; Life Technologies). The stained samples were visualized by using confocal microscopy.

### 2.9 Statistical analysis

Data are presented as mean ± standard error of the mean (SEM). One-way ANOVA analysis was performed to determine statistical significance and data was analyzed for unequal variances using post-hoc tests for multiple comparisons. Results with a p-value less than 0.05 were considered to be significant with *p < 0.05 and **p < 0.01. All analyses were performed using SPSS version 26 (SPSS Inc., USA).

## 3. Results

### 3.1 Proliferation and characteristics of pluripotent ESCs grown in 3-D self-assembling scaffolds

We first investigated the encapsulation and proliferation of Elf1 cells in 3-D self-assembling scaffolds by light microscopy and MTT analysis (Figure 2). The results presented in Figure 2A show that Elf1 cells displayed compact clonal morphology and growth, consistent with naïve ESCs grown in 2-D culture conditions. When Efl1 cells were encapsulated in 3-D scaffolds, they proliferated with a distinctly progressive increase in the size of dark and densely packed colonies (Figure 2B-E). These 3-D grown cells maintained their undifferentiated clonal morphology, indistinguishable from the initial cells, when subcultured back to 2-D culture conditions (Figure 2F). Quantitative analysis showed a steady and significant increase in cell growth over a period of 21 days (Figure 2G). Characterization of 3-D grown Elf1 cells depicted in Figure 2H show upregulation in the expression of core pluripotent markers, *OCT4*, *NANOG*, and *SOX2*, which increased 1.6-, 1.5-, and 2.3-fold, respectively on day 21 of culture. Interestingly, a similar but more pronounced increases were observed in the expression of naïve markers, *KLF17*, *KLF4*, *TFCP2L1*, *DPPA3*, and *DNMT3L*, which increased by 2.6-, 2.2-, 2.7-, 4.4-, and 3.2-fold, respectively. A precipitous decrease in the expression levels of most of the core and naïve genes was observed upon subculturing the 3-D grown Elf1 cells under 2-D culture conditions. The expression levels decreased to levels which were similar to that of the initial 2-D grown cells. Although, the expression of *OCT4*, *DPPA3*, and *DNMT3L* genes remained relatively high, they gradually decreased to normal levels upon repeated passaging under 2-D culture conditions (data not shown). Together these results suggest that the 3-D culture conditions effectively maintained the pluripotent growth of Elf1 cells.

**Figure 2.**
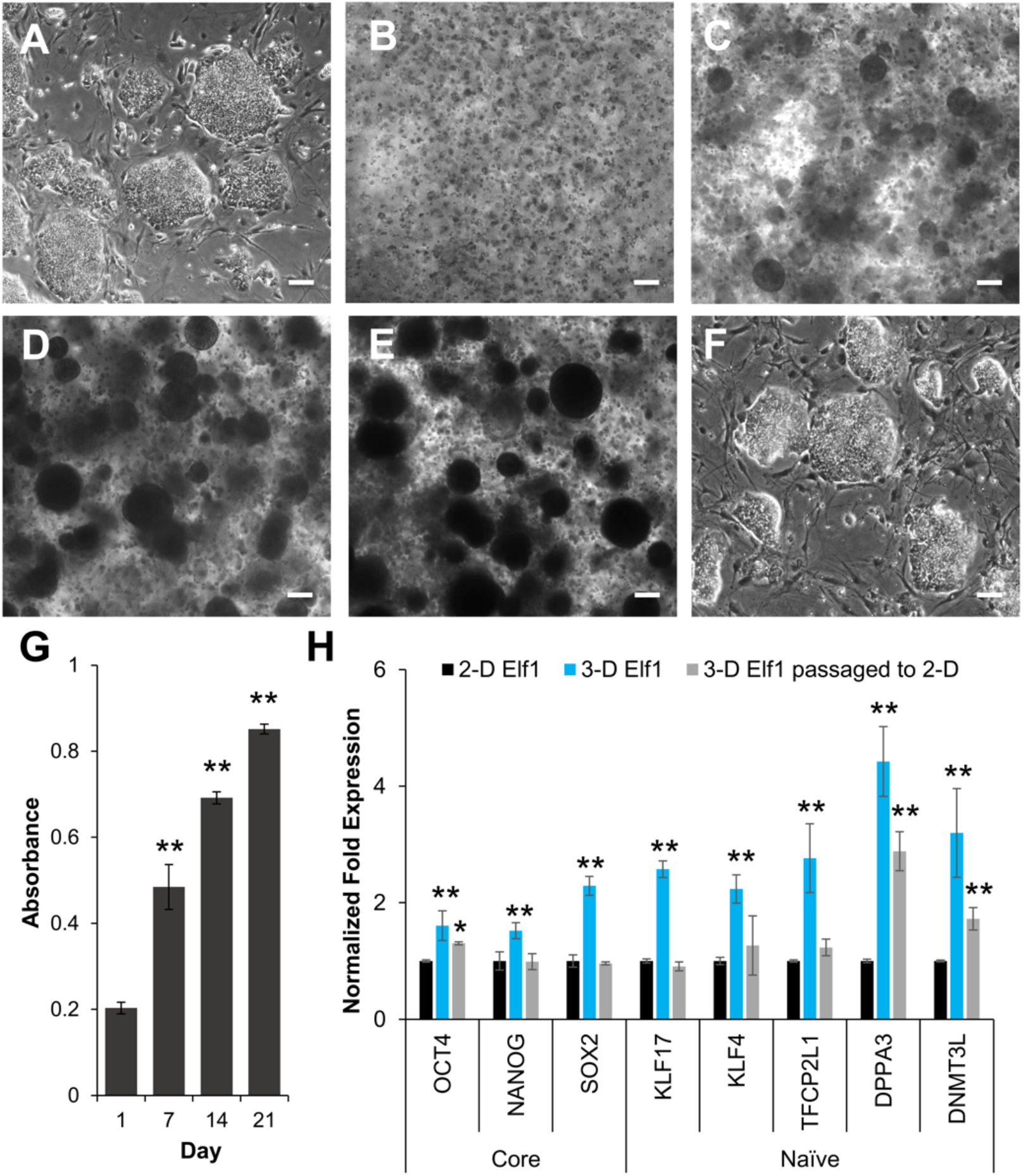
Comparison of naïve ESCs grown in 2-D and 3-D culture conditions. (A) LM showing morphology of Elf1 cells grown in 2-D conditions, (B-E) LM showing morphology of Elf1 cells grown in 3-D PEG-8-SH/PEG-8-Acr scaffolds for 0, 7, 14 and 21 days, respectively. (F) LM of Elf1 cells grown in 3-D and then subcultured back to 2-D culture conditions, displayed undifferentiated morphology and clonal growth comparable to the initial 2-D grown cells. All scale bars represent 100 μm. (G) Quantitative determination of ESC proliferation by MTT assay. Results were expressed as the absorbance ± SEM with a significant increase in cell number. (H) Expression of core and naïve pluripotency markers in Elf1 cells grown in 2-D and 3-D culture conditions as well as 3-D grown cells passaged to 2-D culture as determined by qRT-PCR. Results were expressed as the fold expression ± SE normalized to reference genes *HMBS* and *GAPDH* (*p ≤ 0.05 and **p ≤ 0.01).

These results were then validated by assessing the differentiation potential of 3-D grown Elf1 cells both *in vitro* (data not shown) and *in vivo* by teratoma formation assay. The explanted teratomas were analyzed by immunohistochemical staining and qRT-PCR (Figure 3C and 3D). The results show positive expression of GATA4, BRACHYURY, and TUJ1 proteins, representative of endoderm, mesoderm, and ectoderm tissue, suggesting that 3-D grown Elf1 cells differentiated into all three germ layers *in vivo*. Transcriptional analysis of teratomas also showed expression of germline specific markers, *SOX7* and *SOX17* (endoderm), *MIXL1* and *N-CADHERIN* (mesoderm), as well as *NCAM* and *PAX6* (ectoderm). Overall, these results confirmed that 3-D culture conditions maintained the characteristics of Elf1 cells and upregulated the expression of core naïve genes, suggesting that the self-assembling scaffolds provided a supportive microenvironment for pluripotency.

**Figure 3.**
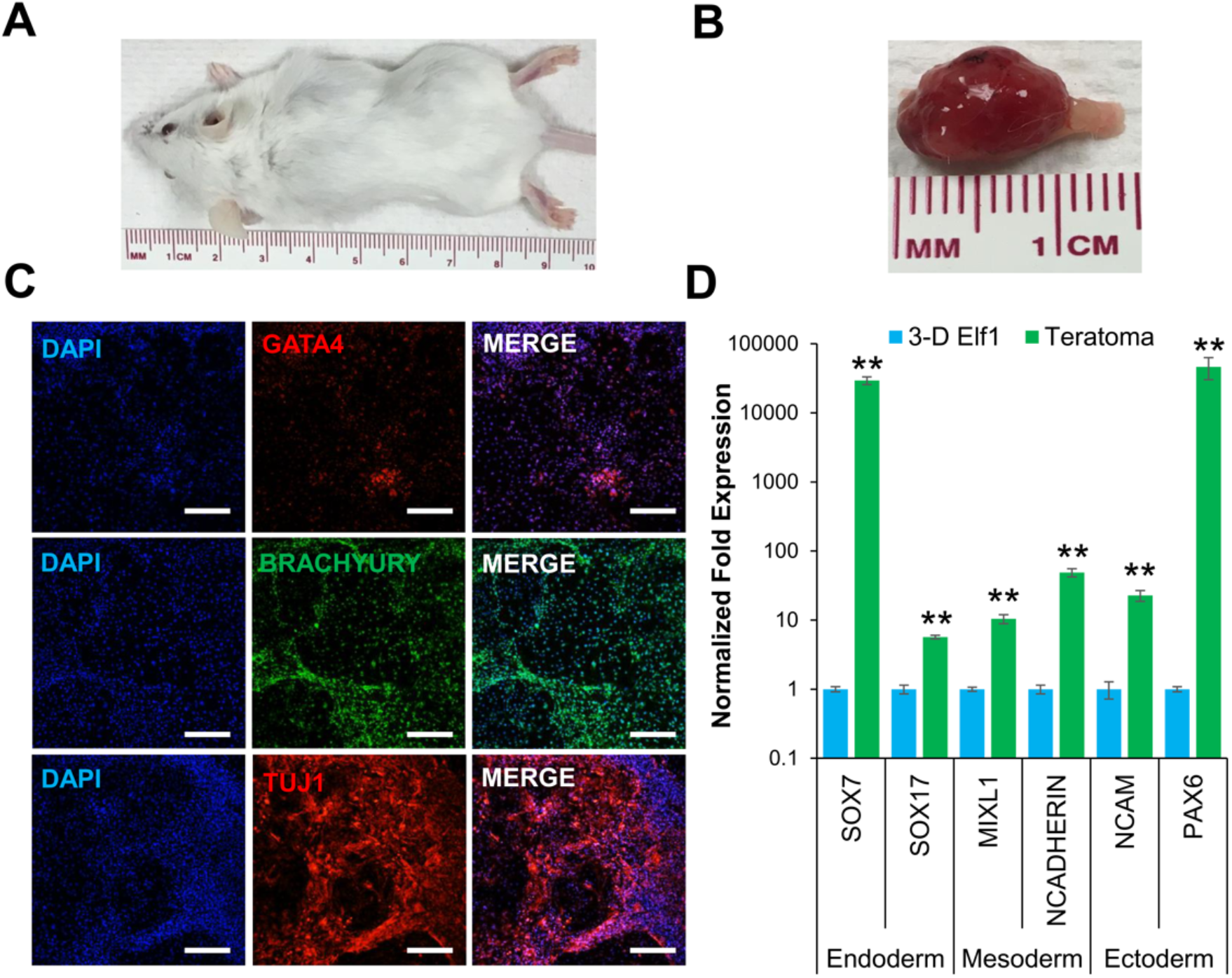
3-D cultured naïve ESCs produced teratomas in SCID-beige mice. (A) Tumor growth was observed in all mice injected with 3-D grown Elf1 cells (*n* = 3). (B) Explanted tumor at 4 weeks showed encapsulated, lobular and well-circumscribed gross morphology consistent with teratoma growth. (C) Confocal images (10X) depict the presence of GATA4, BRACHYURY, and TUJ1 protein expression representing endoderm, mesoderm, and ectoderm, respectively. All scale bars represent 100 μm. (D) Gene expression analysis by qRT-PCR showed that teratomas expressed, *SOX7*, and *SOX17* (endoderm), *MIXL1*, and *N-CADHERIN* (mesoderm), and *NCAM*, and *PAX6* (ectoderm). Results are expressed as the fold expression ± SEM normalized to reference genes *HMBS* and *GAPDH* (*p ≤ 0.05 and **p ≤ 0.01).

### 3.2 Transcriptomic analysis of naïve ESCs grown in 2-D and 3-D culture conditions

Many studies have shown that 3-D cell culture conditions can modulate gene expression via biological and/or mechanical signals [23, 24]. To investigate the molecular level differences in Elf1 grown in 2-D and 3-D culture conditions, RNA-seq was performed. Differential expression analysis of the RNA-seq data identified 802 differentially expressed genes (DEGs) at FDR < 0.01, and 1290 DEGs at FDR < 0.05, with 489 and 792 upregulated genes in 2-D and 3-D grown cells, respectively. The clear distinction of the transcriptomic profile between culture conditions was evident in the heatmap depicting the top 100 DEGs (FDR < 4.0×10^−9^), of which 39 genes were upregulated in 2-D grown cells and 61 genes were upregulated in 3-D grown cells (Figure 4).

**Figure 4.**
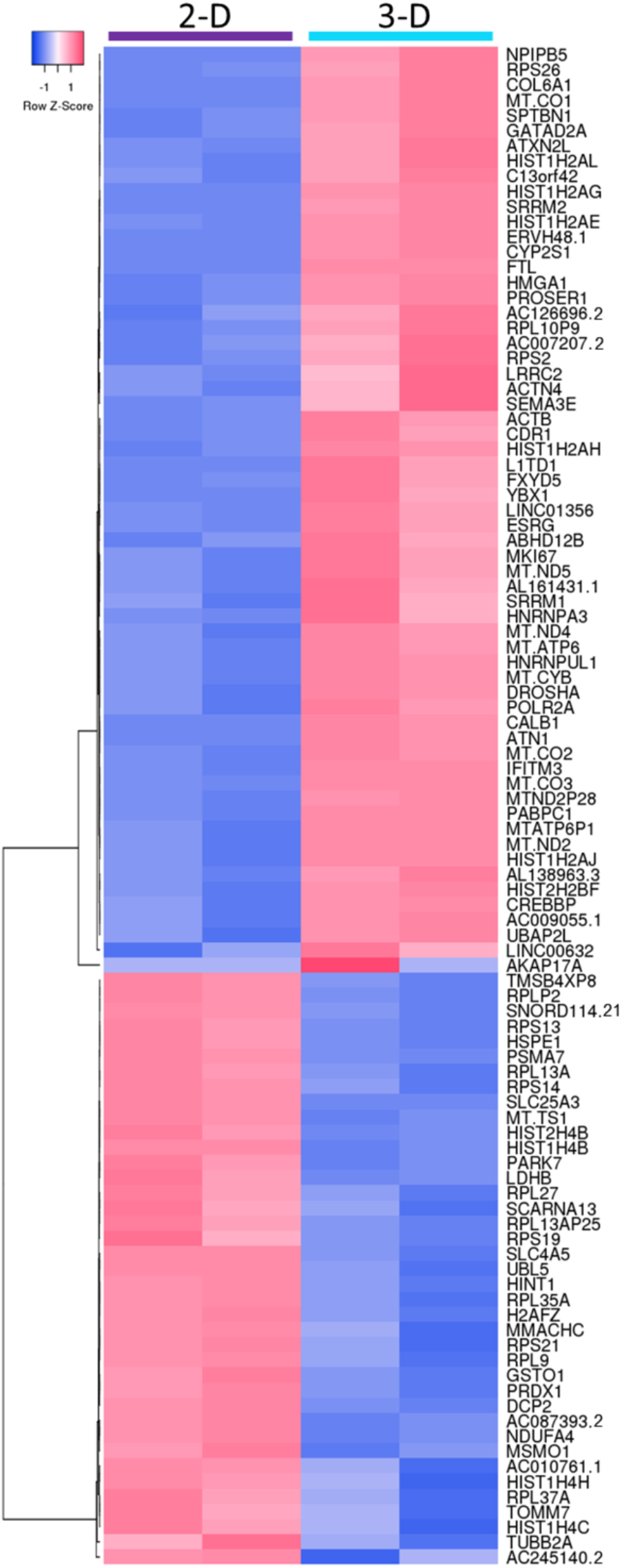
Heatmap showing raw z-scores of RNA-seq log2 transformed values of the top100 differentially expressed genes in 2-D and 3-D cultured Elf1 cells. Up- and down-regulated genes are red and blue, respectively.

### 3.3 Functional analysis of DEGs using gene ontology (GO) and enrichment analysis

GO and pathway analysis of genes with an FDR < 0.05 were performed using Enrichr and PANTHER (Figure 5). Comparative enrichment analysis of Elf1 cells grown in 2-D and 3-D culture conditions performed using Enrichr identified DEGs associated with cellular compartment, molecular function, and biological process ontologies (Figure 5A-C). In the cellular compartment category, 3-D grown cells were found to be highly enriched for GO terms, including focal adhesion (GO:0005925), spliceosomal complex (GO:0005681), stress fiber (GO:0001725), β-catenin-TCF complex (GO:1990907), cytoskeleton (GO:0005856), polymeric cytoskeleton fiber (GO:0099513), chromatin (GO:0000785), microtubule (GO:0005874), histone methyltransferase complex (GO:0035097), and actin cytoskeleton (GO:0015629) (Figure 5A). While upregulated DEGs in 2-D grown cells were only significantly enriched in the cellular compartment categories of focal adhesion and spliceosomal complex, with much lower combined scores when compared to 3-D grown cells.

**Figure 5.**
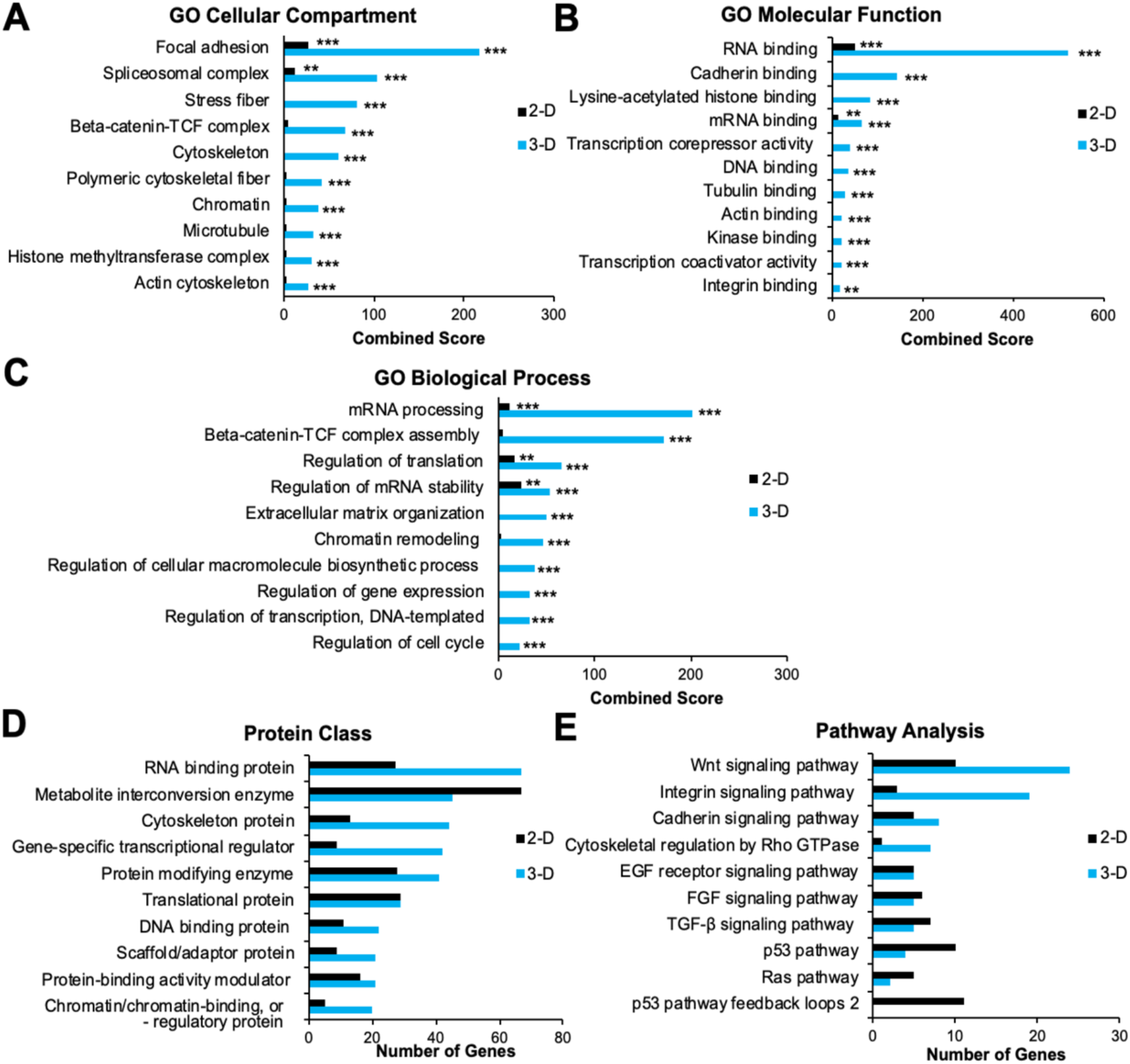
Gene ontology analysis of differentially expressed genes in naïve ESCs cultured under 2-D and 3-D conditions. (A-C) Enrichr and (D-E) PANTHER analysis of differential gene expression with FDR < 0.05 as determined by DESeq2 analysis. (A-C) GO term enrichment analysis depicting upregulated DEGs associated with the cellular compartment, molecular function, and biological processes, respectively. The x-axis represents the combined score (p-value multiplied by the z-score) generated by Enrichr (*p ≤ 0.05, **p ≤ 0.01, and ***p ≤ 0.001). (D-E) Upregulated protein classes and pathways associated with 2-D in 3-D cultured Elf1 cells according to PANTHER analysis, respectively. The x-axis represents the number of genes associated with each category.

GO terms for molecular function of DEGs upregulated in 3-D culture showed significant enrichment in RNA binding (GO:0003723), cadherin binding (GO:0045296), lysine-acetylated histone binding (GO:0070577), mRNA binding (GO:0003729), transcription corepressor binding (GO:0003714), DNA binding (GO:0003677), tubulin binding (GO:0015631), actin binding (GO:0003779), kinase binding (GO:0019900), transcription coactivator activity (GO:0003713), and integrin binding (GO:0005178) (Figure 5B). For these functions, 2-D grown cells showed DEGs in 2-D grown cells were only significantly enriched in the cellular compartment categories of focal adhesion and spliceosomal complex, with much lower combined scores when compared to 3-D grown cells.

GO terms for molecular function of DEGs upregulated in 3-D culture showed significant enrichment in RNA binding (GO:0003723), cadherin binding (GO:0045296), lysine-acetylated histone binding (GO:0070577), mRNA binding (GO:0003729), transcription corepressor binding (GO:0003714), DNA binding (GO:0003677), tubulin binding (GO:0015631), actin binding (GO:0003779), kinase binding (GO:0019900), transcription coactivator activity (GO:0003713), and integrin binding (GO:0005178) (Figure 5B). For these functions, 2-D grown cells showed enrichment of DEGs in categories of RNA binding and mRNA binding at levels significantly lower than that of 3-D grown cells.

When gene products were classified according to biological process GO terms, 3-D grown cells exhibited significant enrichment in pathways/processes associated with mRNA processing (GO:0006397), β-catenin-TCF complex assembly (GO:1904837), regulation of translation (GO:0006417), regulation of mRNA stability (GO:0043488), extracellular matrix organization (GO:0030198), chromatin remodeling (GO:0006338), regulation of cellular macromolecular biosynthetic process (GO:2000112), regulation of gene expression (GO:0010468), regulation of DNA-templated transcription (GO:0006355), and regulation of cell cycle (GO:0051726) (Figure 5C). Of these biological processes, 2-D grown cells displayed significant upregulation of DEGs in mRNA processing, regulation of translation, and regulation of mRNA stability, with slight but not significant increase in β-catenin-TCF complex assembly and chromatin remodeling categories.

GO analysis was also performed using PANTHER to identify protein classes and pathways upregulated in DEGs in 2-D and 3-D culture conditions, with 431 (out of 484) genes and 744 (out of 812) genes mapped to the PANTHER database, respectively. Predominant protein classes expressed in 3-D grown cells include RNA binding protein (PC00031), followed by cytoskeleton protein (PC00085), gene-specific transcriptional regulator (PC00264), protein modifying enzyme (PC00260), DNA binding protein (PC0009), scaffold/adaptor protein (PC00226), protein-binding activity modulator (PC00095), and chromatin/chromatin-binding or -regulatory protein (PC00077) (Figure 5D). On the other hand, a higher number of DEGs were associated with metabolite interconversion enzyme (PC00262) in 2-D cultured cells, and the total number of DEGs associated with translational proteins (PC00263) were the same in cells grown both 2-D and 3-D conditions.

To investigate the effect of 2-D and 3-D culture conditions on cell signaling, pathway analysis of the DEGs was performed. A higher number of genes were associated with Wnt (P00057), integrin (P00034), cytoskeleton regulation by Rho GTPase (P00016), and cadherin (P00012) signaling in 3-D grown cells (Figure 5E). While the number of genes associated with FGF (P00021) and TGF-β (P00052) signaling pathways were slightly higher in 2-D compared to 3-D cultured cells, a greater increase was observed in Ras (P04393), p53 (P00059), and p53 feedback loop (P04398) pathways. In contrast, the same number of genes were differentially expressed in EGF receptor signaling (P00018) in both 2-D and 3-D cultured cells.

These results show enrichment of GO terms were primarily associated with the regulation of gene expression via chromatin, transcriptional, and translational modification as well as organization of the ECM/cytoskeleton. Furthermore, 3-D culture conditions appeared to modulate transcriptional regulation of genes associated with several signaling pathways including Wnt, integrin, cadherin, cytoskeletal regulation by Rho GTPase, and p53.

### 3.4 KEGG pathway analysis of naïve ESCs grown in 2-D and 3-D culture conditions

We then examined the most significantly enriched pathways in cells grown in 2-D and 3-D culture conditions. DEGs were mapped to the KEGG database (p < 0.05; FDR < 0.25) in focal adhesion, Wnt signaling, and p53 signaling pathways and visualized using Venn diagrams and heatmaps (Figure 6). There were 38 genes associated with the focal adhesion pathway upregulated in cells grown in 3-D as opposed to 9 genes in 2-D culture conditions (Figure 6A). Genes upregulated in 3-D grown cells encoded basement membrane proteins (i.e. *COL4A2*, *COL6A1*, *FN1*, *LAMC1*, and *LAMB2*), actin binding proteins (i.e *ACTN4*, *FLNA*, *FLNC*, and *TLN1*), as well as components of the integrin signaling pathway (i.e *CAV1*, *ITGB1*, *ITGA3*, *ITGA5*, and *PXN*), and downstream kinases (i.e *AKT1* and *MAPK3*). Twenty five genes involved in the Wnt signaling pathway were enriched in 3-D grown cells (i.e *TCF7L2*, *TC7L1*, *TCF7*, *CREBBP*, *DVL3*, *RUVBL1*, *SFRP2*, *FOSL1*, *APC*, *AXIN2*, *LEF1*, *LRP5*, and *MYC*) in comparison to only 8 upregulated genes observed in 2-D grown cells (i.e *MAPK10*, *SMAD4*, *FZD7*, and *FZD3*) (Figure 6B). In contrast, upregulation of only 5 genes of the p53 pathway were upregulated in 3-D culture (i.e *THBS1*, *SHISA5*, *SERPINE*, *BCL2L1*, and *SFN*), compared to the 15 genes (i.e *RPRM*, *CCNG1*, *APAF1*, *ATM*, *CHEK2*, *CDK1*, *MDM4*, and *CYCS*) were represented in 2-D grown cells (Figure 6C). Generally, KEGG analysis of enriched pathways suggested that focal adhesion and Wnt pathways were upregulated, while p53 pathway signaling were downregulated upon 3-D culture of Elf1 cells.

**Figure 6.**
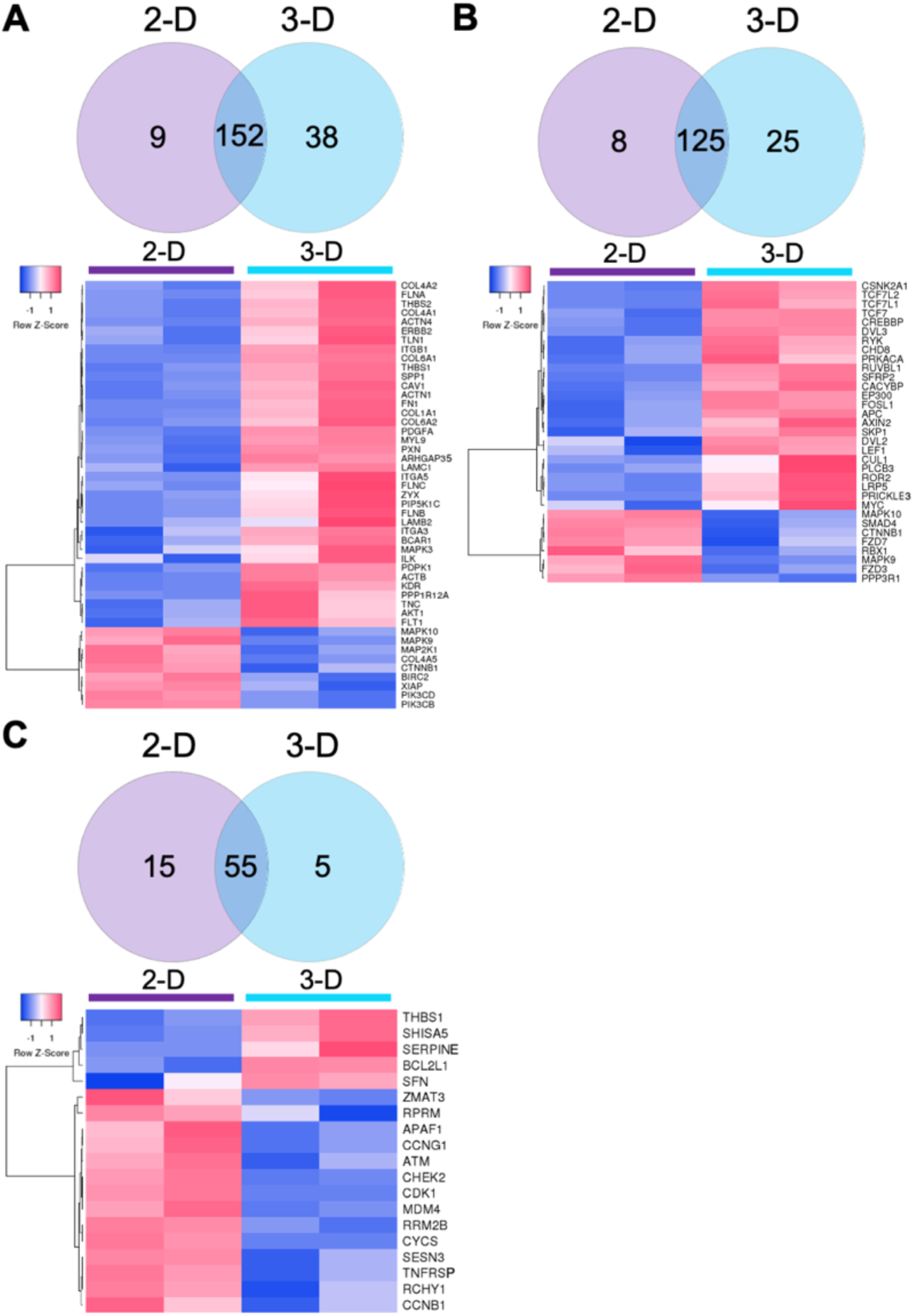
Heatmaps showing raw z-scores of RNA-seq log2 transformed values of 2-D and 3-D grown naïve ESCs. Venn diagrams show the number of genes upregulated in 2-D, genes not significantly changed between 2-D and 3-D, and genes upregulated in Elf1 cells during growth in 3-D scaffolds. Up- and down-regulated genes are red and blue, respectively, at a cutoff of p < 0.05. Heatmaps depicting significantly DEGs associated with (A) the focal adhesion pathway, (B) Wnt pathway, and (C) p53 pathway was determined using KEGG analysis.

### 3.5 Validation of RNA-seq results with qRT-PCR

We next confirmed the RNA-seq data by analyzing selected genes representing various differentially expressed pathways by qRT-PCR (Figure 7). Consistent with the RNA-seq data, the genes associated with focal adhesion including *ITGB1, MYL9, COL6A1*, and *ACTN4* were significantly increased in 3-D relative to 2-D grown cells. Similarly, within the Wnt signaling pathway, *SFRP2*, *RUVBL1, DVL3*, and *CREBBP*, were upregulated and *FZD7* and *SMAD4* were downregulated in cells grown in 3-D compared to 2-D culture conditions. On the other hand, *CDK1*, *MDM4*, and *CCNG1* genes, associated with the p53 pathway, were downregulated in cells grown in 3-D relative to 2-D culture. The trend in the expression of these genes is consistent with what was observed in the RNA-seq data. Taken together, these results lead us to propose a molecular mechanism involved in the self-renewal and pluripotency of Elf1 cells in 3-D culture as depicted in Figure 7B.

**Figure 7.**
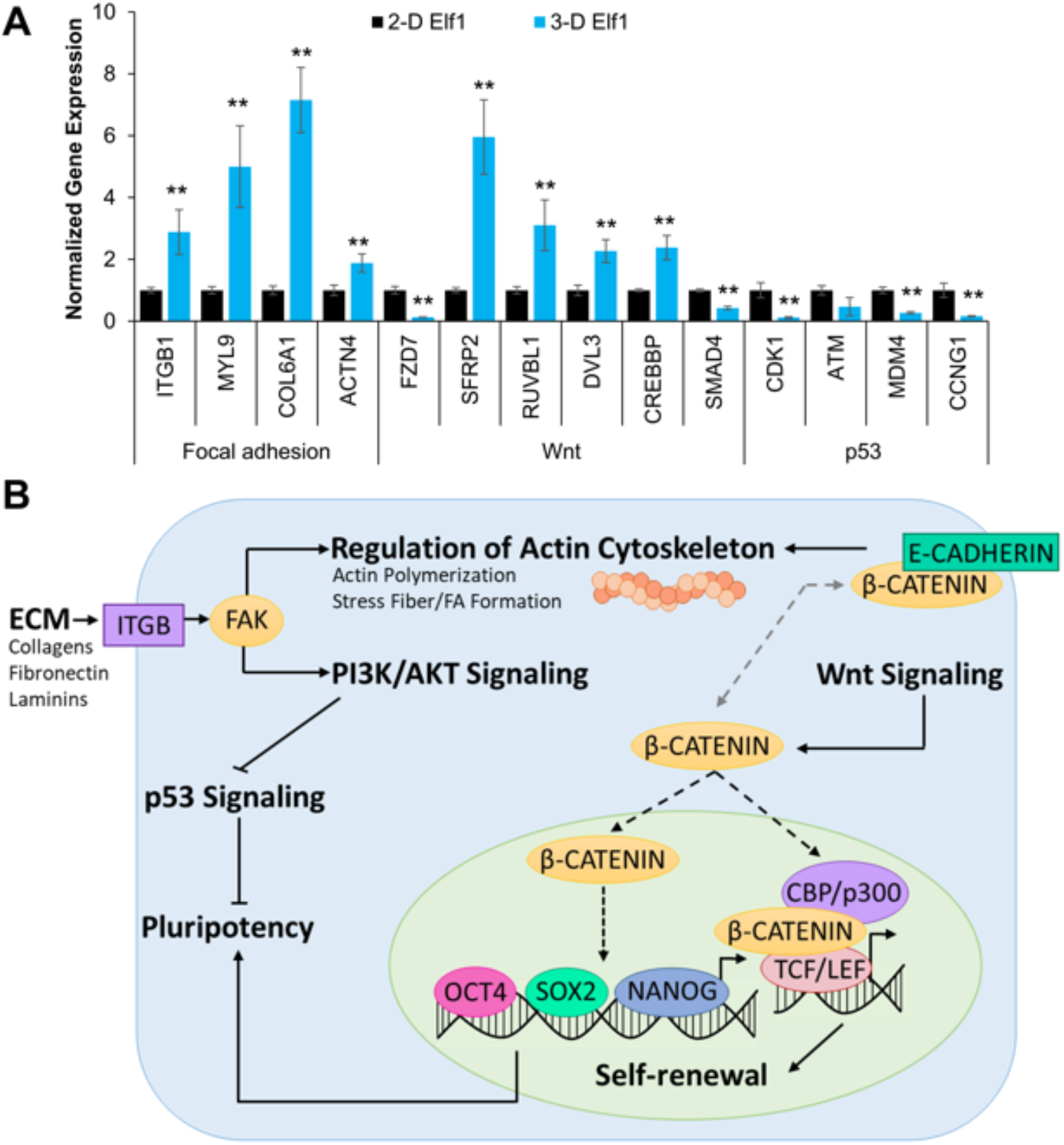
Analysis of genes involved in the maintenance of pluripotency in 3-D grown naïve ESCs. (A) 3-D culture conditions affected the expression of genes associated with focal adhesion, Wnt, and p53 signaling pathways as determined by qRT-PCR. Results were expressed as the fold expression ± SEM normalized to reference genes *HMBS* and *GAPDH* (*p < 0.05 and **p < 0.01). (B) Proposed pathway involved in the maintenance of pluripotency of Elf1 cells grown under 3-D culture conditions.

## 4. Discussion

Two-dimensional culture of human ESCs is marred by batch-to-batch variation, poor viability, and spontaneous differentiation [27, 28]. It is well-recognized that cues from culture conditions including media composition and substrate stiffness influence self-renewal and cell fate [24]. To circumvent these problems, we investigated the effect of 3-D culture on the growth, maintenance, and gene expression of naïve human ESCs.

Elf1 cells grown in scaffolds composed of self-assembling PEG polymers for 3 weeks without passaging maintained their pluripotent properties including self-renewal and differentiation potential as evident by their ability to form teratomas. Growth of cells in the 3-D scaffolds produced darker and more compact colonies compared to cells cultured under 2-D conditions. 3-D grown cells also exhibited increased expression of core as well as naïve pluripotent markers. However, their expression declined to normal levels when subcultured back to 2-D culture conditions, suggesting that 3-D culture conditions including the structure and composition of the scaffolds were responsible for the modulation of gene expression. Previously, maintenance of primed human ESCs in various 3-D culture conditions has been shown to differentially effect *OCT4* expression, which ranged from a slight decrease reported in thermoresponsive synthetic hydrogels [52], to no significant change in natural scaffolds composed of hyaluronic acid [30], chitosan or alginate [53]. However, upregulation of select core pluripotent markers has been observed in 3-D culture of mouse [36, 54] as well as human ESCs [55, 56], suggesting a dependence on scaffold composition and culture conditions. We have reported upregulation of select naïve pluripotent markers in primed [37] and naïve human ESCs during 3-D culture in the PEG-8-SH/PEG-8-Acr scaffolds. To determine the influence of the 3-D culture microenvironment on gene expression profiles in Elf1 cells, we performed global transcriptomic analysis.

We identified a significant number of DEGs in 2-D and 3-D culture conditions. Functional analysis of these genes and overlap of GO terms indicated that 3-D culture conditions impacted regulation of transcription and chromatin as well as the organization of the ECM modulated via various signaling pathways. 3-D grown cells also displayed enrichment of biological process GO terms including mRNA processing, mRNA stability, and translation, which could be responsible for the long-term self-renewal observed upon encapsulation in the self-assembling scaffold. We also observed upregulated GO terms associated with chromatin remodeling and histone methyltransferase activity in 3-D grown naïve ESCs, shown to be involved with increased transcriptional activation [57] and the establishment of induced pluripotency [58]. Upregulation of genes associated with chromatin remodeling in 3-D grown cells may suggest a continued open chromatin structure needed for the genes involved in pluripotency. This would be consistent with the higher expression of many of the observed pluripotent markers. Previous RNA-seq studies comparing naïve and primed human ESC epigenomic dynamics suggest that a more open chromatin structure is associated with naïve pluripotency, with enhancer chromatin and 3-D genome architecture changes reported to be involved in the transition from naïve to primed ESCs [59]. This might imply that the 3-D scaffold environment was more supportive of the maintenance of the naïve pluripotent state.

Extensive overlap of GO terms associated with the organization of the ECM and modulation of the actin cytoskeleton were also significantly overrepresented in 3-D grown cells. 3-D grown cells exhibited increased DEGs encoding major components of the basement membrane synthetized by pluripotent stem cells, including collagen, fibronectin, and laminin, which have been shown to support ESC pluripotency when used as 2-D culture substrates via integrin binding [60]. It is well-known that the transduction of biophysical cues from the microenvironment, including matrix dimensionality, mechanical forces, and cell–cell interactions, can influence ESC shape and fate determination as well as ECM deposition during development [25].

Cell interaction with the surrounding ECM can affect cytoskeletal organization, nuclear morphology, and in turn can affect gene expression via alteration of actin fibers [61]. In our study, 3-D grown cells exhibited clonal growth forming large rounded colonies during growth in the self-assembling scaffold and upregulation of DEGs a sociated with actin cytoskeleton remodeling while maintaining pluripotency. In a recent study, 2-D maintenance of human ESCs using various growth media was shown to differentially influence cytoskeleton protein expression and morphology in a cell line dependent manner, with culture in more defined media resulting in more compact clonal growth [62]. Since, we used the same growth media for both 2-D and 3-D culture, it is plausible that the dimensionality and/or composition of the scaffold was responsible for the observed transcriptional changes associated with ECM and cytoskeletal organization. Furthermore, since the PEG-8-SH/PEG-8-Acr scaffold is biologically inert, the 3-D scaffold provided a more defined microenvironment, which altered mechanical cues typically associated with 2-D culture.

PANTHER pathway analysis of 3-D grown Elf1 cells depicted higher representation of DEGs associated with cytoskeleton regulation by Rho GTPase as well as integrin, cadherin and Wnt signaling; whereas, fewer DEGs were observed in Ras and p53 pathways in comparison to 2-D cultured cells. Based on these findings, we performed further KEGG pathway analyses and proposed a potential mechanism for the long-term maintenance of naïve pluripotency in Elf1 cells grown in the 3-D self-assembling scaffolds.

In both mouse and human ESCs, focal adhesion and integrin signaling serve pleiotropic roles in anchorage, contractility, survival, proliferation as well as support or inhibition of self-renewal [63]. Our study revealed upregulation in 3-D grown cells of several genes such as *FLNC*, *COL4*, *CAV1*, and *SERPINE1*, as well as *ITGB1*, *ILK*, and *AKT1*, which have been reported to be associated with actin cytoskeleton remodeling [64], and integrin mediated focal adhesion activation [65], respectively. Activated integrin signaling stimulates the PI3K/Akt signaling pathway, which results in downstream inhibition of ERK signaling pathway [66], playing an essential role in both mouse and human ESC pluripotency [67]. Additionally, mechanical cues arising from the ECM are transduced to the nucleus through cytoskeleton remodeling and stress fibers via integrins and focal adhesion complexes [68]. Changes in the actin cytoskeleton induced by substrate stiffness are often regulated by activation of Rho GTPases [69] and promote translocation of mechanosensitive transcription factors such as Yes-associated protein (YAP) to the nucleus in human ESCs [70]. Induction of YAP expression was shown to aid in the long-term survival and expansion of human ESCs in 2-D culture [71]. Furthermore, we have shown induction of naïve-like pluripotency in primed human ESCs grown in 3-D culture, which was abolished upon inhibition of YAP [37]. These observations further suggest a mechanistic role of cytoskeleton remodeling and mechanical signaling in the maintenance of pluripotency.

In addition, we observed a differential activation of Wnt signaling in 2-D and 3-D culture conditions at a transcriptional level, which is supported by several studies suggesting that application of mechanical stimuli and alteration in substrate topography can stimulate Wnt signaling, which plays a role in cell differentiation and maturation in adult stem cells [72–74]. In ESCs, Wnt/β-catenin signaling performs functions in self-renewal, differentiation, and lineage commitment [75]. Activation of Wnt signaling has been shown to promote self-renewal of naïve human ESCs, but is not directly required for the expression of pluripotent markers [76]. Upon transition from the naïve to the primed state, Wnt signaling is largely downregulated [77]. Furthermore, sustained activation of Wnt/β-catenin signaling is needed to preserve normal DNA methylation epigenetic stability in mouse ESCs [78]. Our results suggest diverse transcriptional changes associated with the Wnt pathway including upregulation of downstream Wnt targets, including transcription factors, *TCF7*, *TCF7L1*, and *LEF1*, and chromatin remodeling complexes, *CREBBP* and *EP300*, encoding CREB-binding protein (CBP) and p300, respectively. While TCF7 has been shown to be essential for self-renewal in mouse ESCs [79], TCF7L1 plays an important role in human ESC pluripotency via suppression of primitive streak gene expression [80]. Overexpression of TCF7L1 causes downregulation of mesodermal gene expression, and leads to an improved somatic cell reprogramming and colony formation [81]. However, stabilization of β-catenin has been shown to also enhance self-renewal in naïve mouse ESCs in a TCF independent manner, whereby nuclear β-catenin acts by directly interacting with the master transcription factor, OCT4, to induce the expression of pluripotent genes [82]. Coactivators p300/CBP serve redundant roles to stabilize long-range chromatin loops, which aid in the recruitment of other transcription factors and coregulators, thus promoting expression of self-renewal genes [83]. Furthermore, transcriptional coactivator p300 has been shown to directly aid in the recruitment of the NANOG-OCT4-SOX2 complex in mouse ESCs [84].

Interestingly, human ESC self-renewal has also been shown to be enhanced by short-term stimulation of Wnt/β-catenin signaling via upregulation of E-cadherin, and the subsequent activation of the PI3K/Akt pathway [75]. We observed a higher number of upregulated DEGs associated with cadherin signaling in 3-D grown cells. This may suggest a role for the cadherin-mediated cell-cell interactions in the maintenance of the self-renewing state via altered cell-cell interactions in 3-D culture. E-cadherin has been shown to stabilize LIF signaling [85] and control intracellular levels of β-catenin [86]. Furthermore, 2-D grown cells were characterized by upregulated DEGs in the Ras signaling pathway. Ras signaling is repressed in naïve pluripotency and modulates cadherin protein expression and the epigenetic landscape, playing an important role in the transition to primed pluripotency [87]. We observed downregulation of *CTNNB1*, the gene encoding β-catenin, in 3-D grown cells. Since Wnt signaling is regulated by post-translational modifications of β-catenin and extensive cross-talk with other signaling pathways [88], additional studies are warranted to determine the levels of activated β-catenin upon culture in 3-D scaffolds.

Global transcriptome analysis suggested that 3-D culture in the self-assembling scaffold repressed the p53 signaling pathway in Elf1 cells. p53 acts as a transcriptional regulator of cell cycle arrest and apoptosis and is important for the maintenance of genetic integrity in response to various cellular stresses [89]. In ESCs, p53 is present at low levels in the nucleus, and activation is associated with the induction of differentiation, repressing self-renewal via regulation of key pluripotent markers and microRNAs [90–92]. As such, both maintenance of pluripotency and efficient somatic reprogramming require the suppression of p53 signaling [91]. Additionally, integrin-mediation via focal adhesion kinase (FAK) stimulates Akt signaling, resulting in downstream suppression of p53, which promotes cell survival and maintenance of pluripotency [93]. Transcriptional comparison of naïve ESCs showed increased expression of DEGs associated with the p53 pathway and the p53 feedback regulatory loop, involving *APAF1*, *ATM*, and *CYCS*, when cells were cultured in 2-D culture conditions. This may indicate that 2-D culture resulted in increased heterogeneity due to spontaneous differentiation [27] or enhanced proliferation [94], well-known limitations of traditional planar culture [6].

Overall, 3-D culture conditions appeared to modulate transcriptional regulation of several signaling pathways associated with adhesion and self-renewal, implicating that transduction of mechanical cues from the surrounding scaffold microenvironment may play a role in the regulation of self-renewal and pluripotent marker expression in 3-D culture conditions.

Although these results based on global gene expression are interesting, further analyses are needed to delineate specific mechanistic effects of 3-D culture at a translational level. Nevertheless, these findings suggest the importance of mechanical signaling and culture dimensionality on ESC self-renewal and may foster further developments in long-term 3-D culture of ESCs and their use for cell therapy and regenerative medicine.

## Abbreviations

2-D: two-dimensional
3-D: three-dimensional
CBP: CREB-binding protein
DEGs: differentially expressed genes
ECM: extracellular matrix
EpiSCs: epiblast stem cells
ESCs: Embryonic stem cells
FAK: focal adhesion kinase
FDR: false discovery rate
GO: Gene Ontology
KEGG: Kyoto Encyclopedia of Genes and Genomes
MEF: mouse embryonic feeder
PANTHER: Protein ANalysis THrough Evolutionary Relationships
PEG: polyethylene glycol
PEG-8-Acr: 8-arm PEG acrylate
PEG-8-SH: 8-arm PEG thiol
qRT-PCR: quantitative real time-polymerase chain reaction
RIN: RNA integrity number
RNA-seq: RNA-sequencing
SCID: Severe combined immunodeficiency
SEM: standard error of the mean
YAP: Yes-associated protein

## Acknowledgments

The study was supported by the OU-WB Institute for Stem Cell and Regenerative Medicine (ISCRM), and Oakland University. C. McKee received the Provost Graduate Research Award from Oakland University for this project.

## Data availability

The raw/processed data required to reproduce these findings cannot be shared at this time due to technical or time limitations.

